# A unified framework for dissecting the effects of common signals on functional and effective connectivity analyses: power, coherence, and Granger causality

**DOI:** 10.1101/186122

**Authors:** Dror Cohen, Naotsugu Tsuchiya

## Abstract

When analyzing neural data it is important to consider the limitations of the particular experimental setup. An enduring issue in the context of electrophysiology is the presence of common signals. For example a non-silent reference electrode adds a common signal across all recorded data and this adversely affects functional and effective connectivity analysis. To address the common signals problem, a number of methods have been proposed, but relatively few detailed investigations have been carried out. We address this gap by analyzing local field potentials recorded from the small brains of fruit flies. We conduct our analysis following a solid mathematical framework that allows us to make precise predictions regarding the nature of the common signals. We demonstrate how a framework that jointly analyzes power, coherence and quantities from the Granger causality framework allows us to detect and assess the nature of the common signals. Our analysis revealed substantial common signals in our data, in part due to a non-silent reference electrode. We further show that subtracting spatially adjacent signals (bipolar rereferencing) largely removes the effects of the common signals. However, in some special cases this operation itself introduces a common signal. The mathematical framework and analysis pipeline we present can readily be used by others to detect and assess the nature of the common signals in their data, thereby reducing the chance of misinterpreting the results of functional and effective connectivity analysis.

## Introduction

Understanding how brain areas communicate is one of the fundamental goals of neuroscience. Such neuroscientific investigations often examine communications between brain areas by assessing correlational relationships, known as functional connectivity, or causal relationships, known as effective connectivity (1). Functional and effective connectivity are thought to be involved in fundamental processes such as attention (2-4) and arousal (5-8) and may also be altered in several brain disorders (9). Such connectivity analysis may focus on different spatiotemporal scales, from brain-wide connectivity in the order of seconds, as revealed by functional magnetic resonance imaging (10), to micrometer and millisecond resolution in cortical laminar recordings (11, 12).

However, methodological issues regarding the estimation of functional and effective connectivity are a matter of continuing debate (see (13, 14) for a recent example). An enduring issue in functional and effective connectivity analysis is the adverse effect of common signals in the data. In the context of electrophysiology, a common signal maybe introduced by a non-silent reference electrode or volume conduction/field spread. In both cases the common signal can substantially alter the results of functional or effective connectivity analysis (15-19). To this end, numerous techniques of varying complexity have been suggested for reducing the effect of the common signals ((16, 20-28) and others). One of the simplest and most commonly used techniques is to simply subtract nearby signals, known as bipolar rereferencing.

Even though the adverse effects of common signals on functional and effective connectivity analysis are well recognized, relatively few studies have investigated these effects in depth. In a recent effort to address this, Shirhatti et al investigated how different rereferencing techniques (including bipolar rereferencing) affect the estimation of different neurophysiological metrics using local field potentials (LFPs) recorded from the visual areas of monkeys (29). They found that a measure of functional connectivity known as phase coherence was substantially affected by the choice of the rereferencing scheme. In particular, they claimed that bipolar rereferencing can result in artefactually high phase coherence for specific signal pairs. On the other hand, with respect to effective connectivity, Trongnetrpunya et al used simulations and analysis of LFPs from rats, monkey and human to claim that bipolar rereferencing is effective at removing the adverse effect of common signals for Granger causality analysis (17). The issue of which rereferencing scheme is advantageous and disadvantageous in distinct analysis methods remains an open question.

To date, the effects of common signals on functional and effective connectivity have not been studied in a unified manner. This may be a significant oversight because coherence and Granger causality, two of the most common functional and effective connectivity metrics, are in fact analytically related (30, 31). Analyzing coherence and Granger causality together may help detect common signals, and can also be used to quantify the extent to which various techniques, such as bipolar rereferencing, can remove their adverse effects.

Here, we use LFPs recorded from the brains of fruit flies to analyze power, coherence and Granger causality and demonstrate how these can be jointly used to detect and assess common signals. We conduct our analysis alongside a solid mathematical framework and provide illustrative simulations to highlight the important points. We show that in our data substantial common signals are present, in part due to a non-silent reference signal. Our analysis further suggests that bipolar rereferencing largely removes the effects of the common signals. However, in special cases bipolar rereferencing actually introduces a common signal. The analysis framework we present can be used by others to 1) detect common signals and 2) assess different methods for mitigating their adverse effects.

The paper is structured as follows. The Results section is divided into three subsections corresponding to the analysis of power, coherence and Granger causality. In each section we begin by presenting the relevant theoretical background. The theoretical concepts are complemented by illustrative examples or simulations. Each subsection concludes with the experimental results from the analysis of the fruit fly data. Further methodological details are presented in the accompanying Methods section.

## Results

The theoretical results we present for power and coherence follow from linear dynamics (e.g. (32)) and have been reported elsewhere (e.g. (16-20, 23)). The theoretical results for Granger causality analysis can be found in (30, 31, 33, 34). Our purpose here is 1) to present these results in a single framework, 2) to illustrate the theoretical concepts using simple simulations, and 3) to demonstrate how these relate to analysis of empirical data.

## 1 Power

### 1.1 Theoretical background

#### 1.1.1 Unipolar signal power

We consider the general case in which the signal recorded by some electrode i at time t *y*_*i*_(*t*) represents the sum of a neural signal of interest near the electrode *x*_*i*_(*t*) and a common signal *u*(*t*) that is added to all electrodes

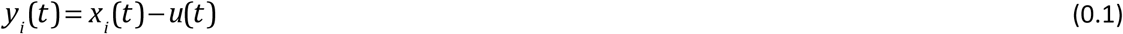

This framework has been used extensively to investigate the case where the common signal corresponds to a non-silent reference signal (e.g. (16, 17, 19, 20, 23)). We refer to *y*_*i*_(*t*) as a unipolar signal.

In the frequency domain Eq. (0.1) becomes

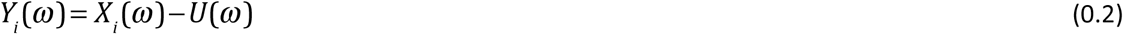

where *Y*_*i*_(*ω*), *X*_*i*_(*ω*) and *U*(*ω*) are the Fourier transforms of *y*_*i*_.(*t*), *x*_*i*_(*t*) and *u*(*t*) respectively.

The power of the unipolar signal is given by

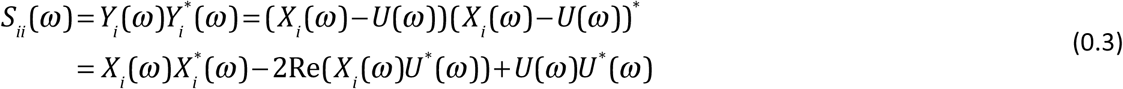

where Re(*X*_*i*_(*ω*)*U*^*^(*ω*)) denotes taking the real part of the cross-spectrum, also known as the co-spectrum. The co-spectrum captures the effect of zero-lag correlations on the power spectrum. If the common signal is uncorrelated with the neural activity (as may be expected from a noisy reference signal) then the co-spectrum vanishes and the power of the unipolar signal is the sum of the power of the neural and common signals

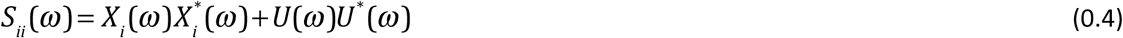

In this case the power of the unipolar signal is an over-estimate of the power of the neural signal.

#### 1.1.2 Bipolar signal power

The experimenter typically only has access to the unipolar signal *y*_*i*_(*t*), not directly to the neural signal *x*_*i*_(*t*) or the common signal *u*(*t*). The challenge is thus to try and remove, or at least reduce, the contribution of the common signal.

When unipolar activity is simultaneously recorded at two nearby locations one can use the additional signal to reduce the effect of the common signal. The simplest strategy is to take the difference between unipolar signals recorded by nearby electrodes. Specifically, given two nearby unipolar signals *y*_*i*_(*t*) and *y*_*i*+1_(*t*) the bipolar rereferenced signal *by*_*i*_(*t*) is defined as

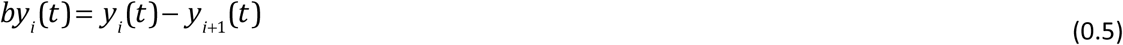

In many recording setups two nearby electrodes may be considered to equally reflect the common signal. If the contribution of the common signal to nearby unipolar signals is identical, then

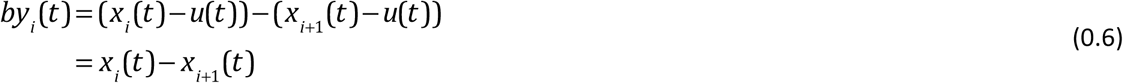

Thus, bipolar rereferencing can provide an estimate of local neural activity that is free from the effect of the common signal.

In the frequency domain Eq. (0.6) becomes

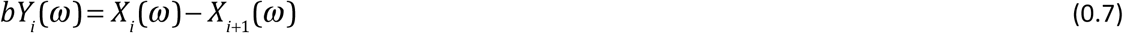

The power of the bipolar signal is given by

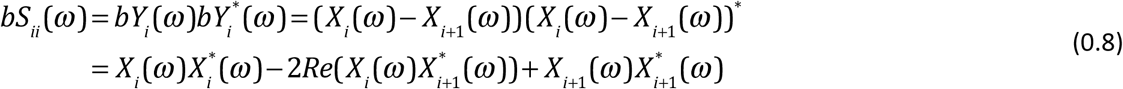

It is useful to compare the expression for the unipolar (Eq. (0.3)) and bipolar signal power (Eq. (0.8)). In general, to completely characterize the unipolar (*S*_*ii*_(*ω*)) and bipolar (*bS*_*ii*_(*ω*)) power, we would need to know 1) the power of neural signals 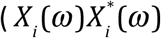 and 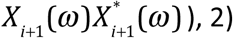 the power of the common signal (*U*(*ω*)*U*^*^(*ω*)) and 3) the relevant co-spectra 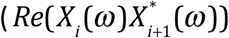 and *Re*(*X*_*i*_ (*ω*)*U*^*^(*ω*))). For example, assuming that co-spectra are negligible in both cases, a stronger common signal (*U*(*ω*)*U*(*ω*)) than the power of neural signals 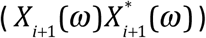 can result in greater unipolar than bipolar power. Another interesting case is when the neural signals recorded by nearby electrodes are very similar, (i.e., *X*_*i*_(*ω*)~ *X*_*i*+1_(*ω*)). In this case, the bipolar power is obviously close to zero, while the unipolar power can be much greater than zero.

In the context of neurophysiology, both scenarios are possible (e.g., substantial non-silent reference, highly similar recordings by nearby electrodes). Under these circumstances we would expect the unipolar signal power to be greater than the bipolar signal power. Theoretically speaking, however, it is entirely possible for the bipolar power to be *greater* than the unipolar power. For example if the power of the neural signals is equal, 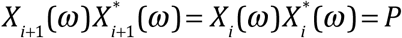 and if the neural signals are independent 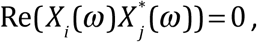 then the power of the bipolar signal is *bS*_*ii*_(*ω*) = 2*P*. If the common signal is comparatively small, e.g. *U*(*ω*)*U*^*^(*ω*) = *P*/2, then the power of the unipolar signal is 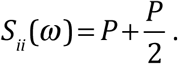.

### 1.2 Experiment

#### 1.2.1 Unipolar power is greater than bipolar power

The mathematical framework we presented demonstrates that in the presence of a common signal the unipolar signal power can be either greater or lesser than the bipolar signal power. Next we empirically quantified unipolar and bipolar power for LFPs recorded with a micro-electrode linear array inserted in the brains of behaving fruit flies (Figure 1, see Methods for details). We considered signals recorded with respect to a reference electrode inserted in the body (thorax) of the fly as unipolar (Figure 1b). We derived bipolar signals by subtracting adjacent unipolar signals (Figure 1c, see Methods for details). In our data we expected unipolar to be greater than bipolar power because 1) the reference signal in the thorax is unlikely to be completely silent and 2) neighboring electrodes are likely to reflect highly similar neural activity due to the close proximity of the electrodes (i.e., 25μm). Consistent with this we found that unipolar was indeed greater than the bipolar signal power (Figure 1d).

**Figure 1.**
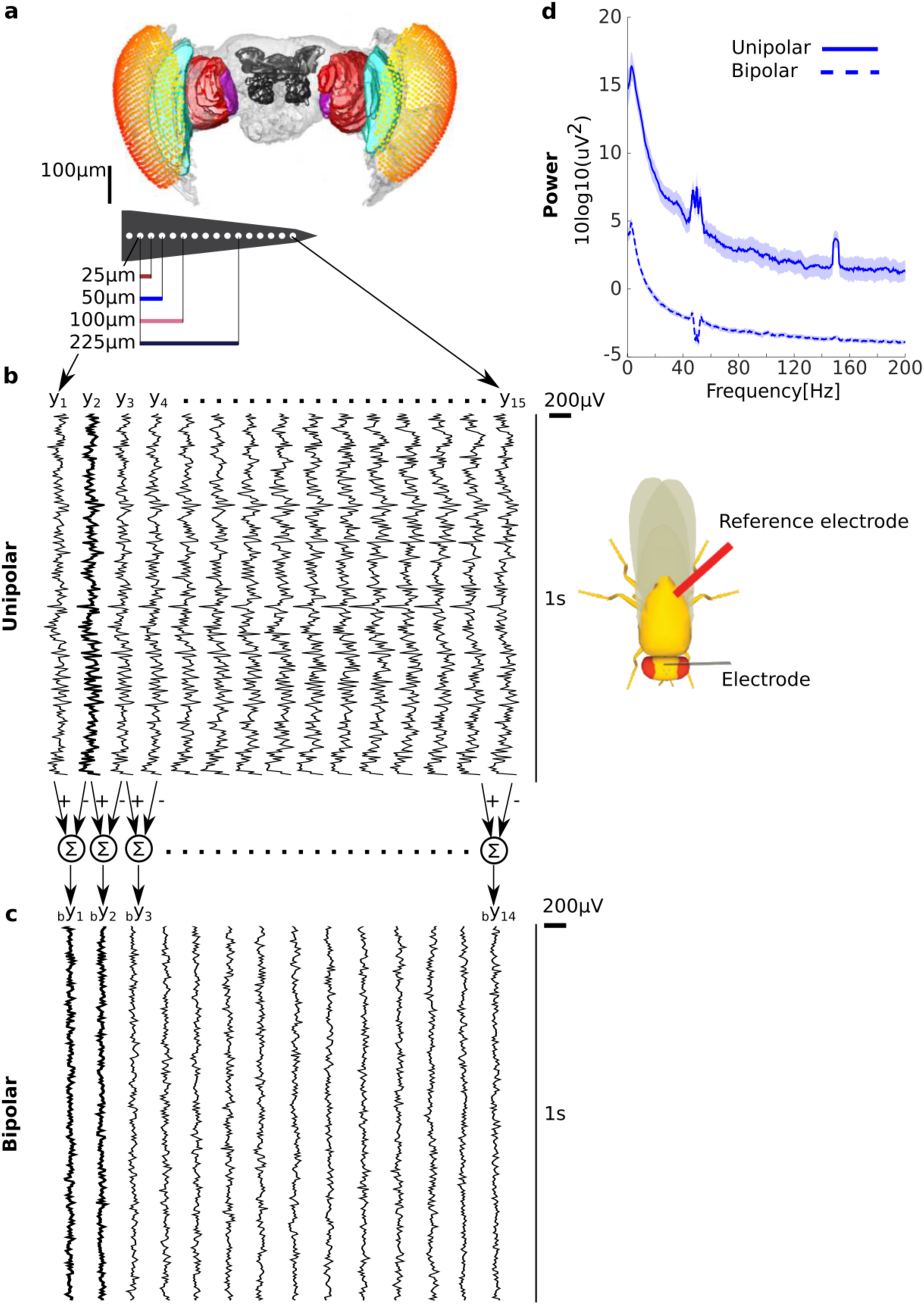
Unipolar power is greater than bipolar power in fruit fly LFP. **a**) Microelectrode recordings of LFPs from fruit flies. Adjacent electrodes are separated by 25μm. **b**) Example of unipolar signals from one fly. The unipolar signals were recorded with reference to the thorax (location shown on schematic on the right). The example clearly shows that the signals are highly correlated across electrodes but that any neighboring pairs are not the same. **c**) Example of bipolar signals derived from two adjacent unipolar signals. The bipolar signals are smaller in magnitude and appear less correlated than the unipolar signals. Two bipolar signals (by1 and by2) that share a unipolar signal (y2 in **b**) in their derivation are shown in bold. **d**) Group average unipolar (solid) and bipolar (dashed) power (N=13, shading reflects standard error of the mean across flies). The much greater unipolar than bipolar power is consistent with a substantial common signal and the highly similar neural activity of adjacent unipolar signals. Peaks at 50 and 150Hz reflect line noise and its harmonic.

## 2. Coherence

### 2.1 Theoretical framework

#### 2.1.1 Unipolar signal coherence

Coherence between signals *Y*_*i*_(*ω*) and *Y*_*j*_(*ω*) measures the extent of linear dependency at each frequency (32) and is defined as:

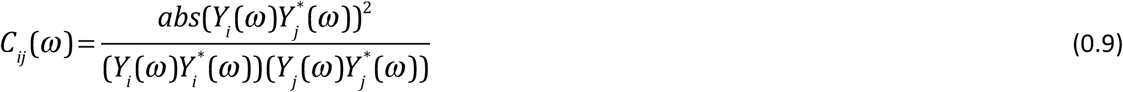

As we will show, in the presence of a common signal, coherence can be non-zero even when the neurophysiological signals (*X*_*i*_(*ω*) and *X*_*j*_(*ω*)) themselves are independent. To see the effect of a common signal on unipolar signal coherence we substitute Eq. (0.2) into (0.9), obtaining:

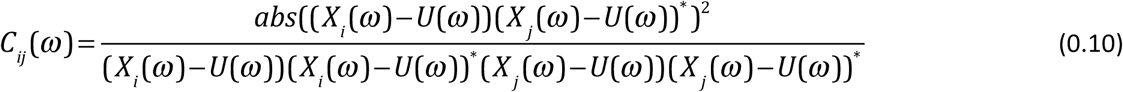

If the common signal and neural signals are independent then the cross-spectrum between the two vanishes and Eq. (0.10) becomes

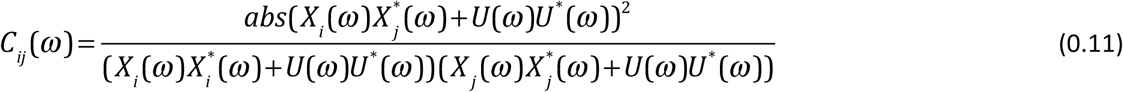

Because we do not have direct access to each of the individual quantities we cannot in general assess the contribution of the common signal to coherence. Here we suggest using prior knowledge of the system in question to try and isolate the relative contribution of the common signal. For example, for neural systems we would not expect genuine neurophysiological coupling for very high frequencies (especially when electrode i and j are sufficiently distant). That is, for high frequencies *ω*_*h*_ we assume that 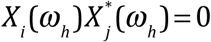. In this case Eq. (0.11) becomes

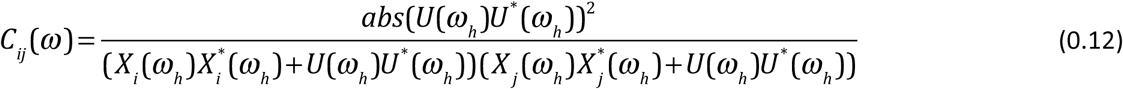

This shows that a common signal can render coherence non-zero even when the neurophysiological sources themselves are not coherent at all. Note that what constitutes such “high frequencies” depends on the system at question. For example coherence at frequencies >100Hz has been reported in mammals (see (35) for discussion of high frequency coherence). Whether these reports were affected by the presence of a common signal is unknown. To date, coherence above 100Hz has not been reported in flies.

Equation (0.12) can be further simplified. First, we define the Neural to Common signal Ratio as 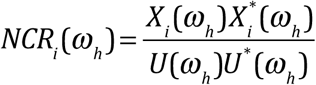

Second, assuming that the NCR is comparable across all electrodes (*NCR*_*i*_(*ω*_*h*_)~*NCR*_*j*_(*ω*_*h*_) = *NCR*(*ω*_*h*_)) we obtain coherence as a function of a single parameter, *NCR*(*ω*_*h*_),

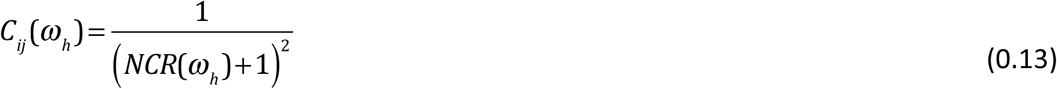

Using Eq. (0.13) we can get an estimate of the NCR only using the observed coherence values

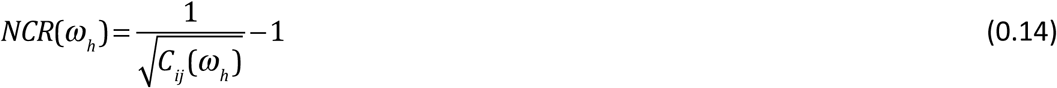

So far the theoretical framework assumes that a single common signal u(t) enters all unipolar signals equally. We note that in real data it is possible that unipolar signals recorded by electrodes that are spatially nearby are affected by multiple common signals. This may be the case if unipolar signals recorded from nearby electrodes pick up the activity of a shared neural source, potentially due to volume conduction/field spread effect (15), in addition to the non-silent reference signal. Unipolar signals recorded by electrodes that are far apart are less likely to reflect neural activity from a shared neural source but may still be affected by the non-silent reference signal.

#### 2.1.2 Bipolar signal coherence

We can investigate the effects of bipolar rereferencing on coherence in a similar way. Substituting Eq. (0.7) into Eq. (0.9) we get the following expression for bipolar signal coherence *bC*_*ii*_(*ω*)

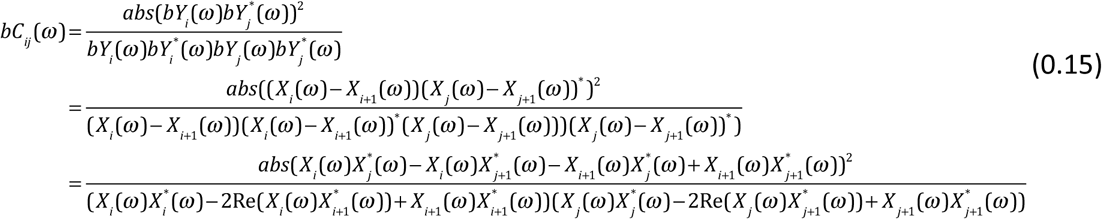

Note that if all combinations of the neural signals are independent (i.e. when i~=j,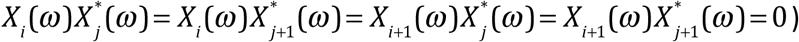 then the cross-spectra in the numerator all vanish and coherence will equal zero. This is in contrast to unipolar coherence, which can be above zero even if the neural signals are independent (Eq. (0.12)).

Coherence between bipolar signals that are derived from a shared unipolar signal (Figure 1c) *bC*_*ii*+1_(*ω*) constitutes a special case,

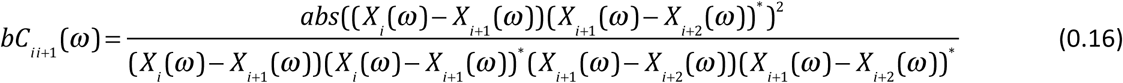

By noting that coherence is invariant with respect to scalar multiplication of the signals, we can use *bY*_*i*+1_(*ω*)→–*bY*_*i*+1_(*ω*) = *X* (*ω*)–*X*_*i*+1_(*ω*). Eq. (0.16) becomes

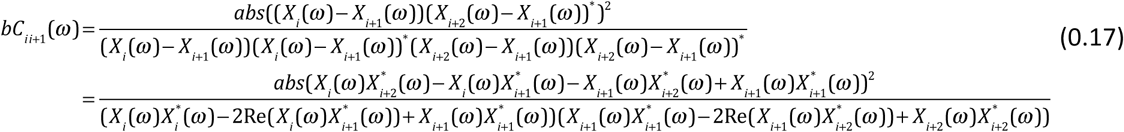

This expression is identical to the case of unipolar coherence (Eq. (0.10)) where the common signal *U*(*ω*) has been replaced with the neural signal from the shared unipolar signal *X*_*i*+1_(*ω*). Intuitively, the shared neural signal acts as a common signal.

If the signals *X*_*i*_(*ω*_*h*_), *X*_*i*+1_(*ω*_*h*_) and *X*_*i*+2_(*ω*_*h*_) are independent for high frequencies *ω*_*h*_, then

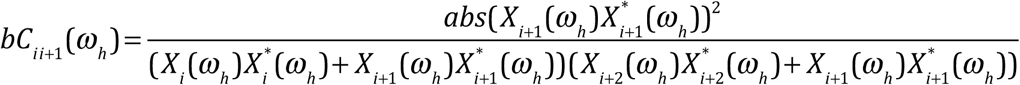

Thus, coherence between bipolar signals that share a unipolar signal in their derivation can be above zero even if neural activity is independent. For example, if the power of the neural activity is equal across channels i, i+1 and i+2 (i.e.,

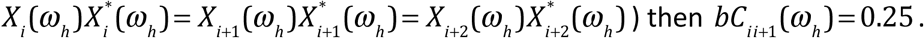

### 2.2 Example simulations: The effect of a common signal on a unidirectionally-connected and a disconnected system

The theoretical framework describes how a common signal can affect coherence. We next use simple simulations to illustrate how the presence of a common signal may manifest in the analysis of real data. To do this we considered four scenarios (Figure 2a-d). Scenarios 1 and 2 correspond to a unidirectionally-connected and a disconnected neural system (meaning that the cross-spectrum between the signals is zero for all frequencies). These two scenarios represent the ‘ground truths’. Scenarios 3 and 4 represent the same two systems but in the presence of common signal that is uncorrelated with both components of the system.

**Figure 2.**
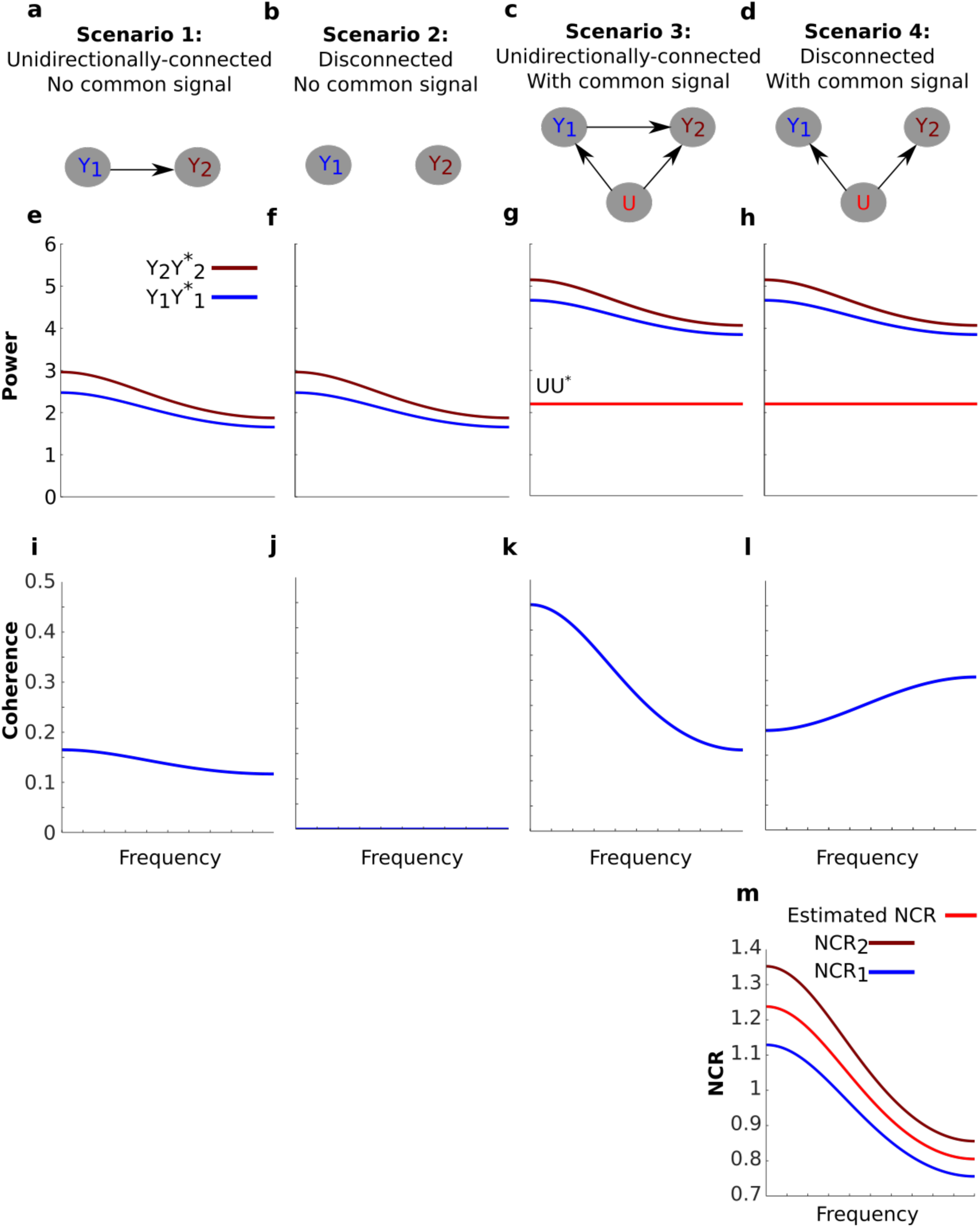
The effect of a common signal on coherence is highly dependent on the spectral characteristics of the system. (**a-d**) Schematics of four simulated scenarios corresponding to a unidirectionally-connected system without a common signal (**a**), a disconnected system without a common signal (**b**), a unidirectionally-connected system with a common signal (**c**) and a disconnected system with a common signal (**d**). (**e-h**) the power of the signals for each scenario (arbitrary units). The systems were constructed to have identical power spectra (**e-f**). The effect of the common signal on power is an increase equal to the power of the common signal (UU*, horizontal red line). The effect of the common signal on power is identical for both systems (**g-h**). (**i-l**) Coherence between the signals for each scenario. Coherence for the unidirectionally-connected system is low and decays with frequency (**i**) and coherence for the disconnected system is zero (**j**). When a common signal is added to the unidirectionally-connected system coherence values increase overall but still decay with frequency (**k**). When a common signal is introduced to the disconnected system coherence values also increase, but now coherence increases with frequency (**l**). This unexpected increase is explained by the Neural to Common signal Ratio (NCR) of this particular system (Eq. (0.13)). (**m**) NCR1 (blue) and NCR2 (brown) refer to the NCR of nodes 1 and two respectively. Because the NCRs for this system decrease with frequency, coherence increases with frequency. We also provide the NCR as estimated directly from coherence (red, Eq. (0.14)).

We modeled the unidirectionally-connected system as a bivariate autoregressive process (see Methods for details)

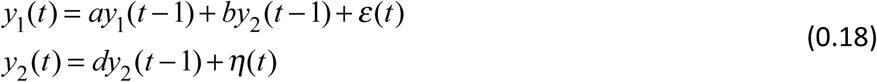

where a, b and d are the autoregressive coefficients and *ε*(*t*) and *η*(*t*) represent uncorrelated noise sources. For autoregressive systems such as this, power, coherence, and Granger causality can all be directly calculated from the autoregressive parameters (see Methods for details).

We set the parameters of the autoregressive process such that the power spectra of the unidirectionally connected system decay with frequency, roughly reflecting a biological system (Figure 2e). We intentionally constructed the disconnected system such that its power spectrum is identical to the unidirectionally-connected system (Figure 2f, see Methods for details). Thus the two systems are indistinguishable based on their power spectra alone. The effect of the uncorrelated common signal on power is given by Eq. (0.4). This effect corresponds to an increase in power equal to the power of the common signal. Note that Eq. (0.4) is independent of the connectivity of the system, so the effect of the common signal on power is identical for the unidirectionally-connected (Figure 2g) and disconnected systems (Figure 2h). Thus, power analysis alone cannot distinguish the two systems.

Next we assessed coherence. The parameters of the autoregressive process result in low coherence values that decay with frequency for the unidirectionally-connected system (Figure 2i). For the disconnected system coherence is obviously zero (Figure 2j). The predominant effect of the common signal on coherence for the unidirectionally-connected system is an overall increase (Figure 2k). Thus, this scenario demonstrates how the presence of a common signal can lead to an overestimation of coherence. For the disconnected system (Figure 2l), coherence values are also increased (from zero), but result in a coherence spectrum that *increases* with frequency.

Though the increase in coherence with frequency is surprising, the theoretical framework nonetheless explains it. Specifically, for the disconnected system coherence is inversely related to Neural Signal to Common signal Ratios (NCRs, Figure 2m, Eq. (0.13)). Because the NCRs of both signals decrease with frequency, coherence increases with frequency. In fact, if we assume that the NCR of both signals are identical then we can *use* coherence to estimate the NCR (Eq. (0.14)). Because for this example the NCRs of both signals are similar, the assumption roughly holds and the resulting estimate is meaningful.

We emphasize that the power spectra of the neural signals and the common signal are identical for scenarios 3 and 4. The only difference is that the cross-spectrum is non-zero for the unidirectionally-connected system, and so the analytical expression for coherence is given by Eq. (0.11). For the disconnected system the cross-spectrum is zero, so coherence is given by Eq. (0.12). In sum, our simulations demonstrate how the mathematical framework we presented may manifest in empirical analysis and illustrate that the effects of a common signal on coherence can be highly dependent on the specifics of the system in question.

### 2.3 Experiment

#### 2.3.1 LFPs recorded in flies show very high coherence with unipolar referencing

Based on the theoretical framework and simulations (Figure 2), we expect that coherence between unipolar signals to present the following characteristics. First, when they are spatially close (25μm), they would be influenced by a stronger common signal than those pairs that are spatially far apart (due to field spread, decay of electrical signal with to the distance). Second, regardless of the distance between the signal pairs, we do not expect to find significant coherence in high frequencies (to our best knowledge, coherence for LFPs in flies has not been reported for frequencies greater than 100Hz (36-39)). Conversely, high coherence in high frequencies indicates the presence of common signals.

We analyzed coherence as a function of the distance between the unipolar signals. To do this we averaged coherence over unipolar signal pairs separated by 25,50,75-150 and 175-325μm (Figure 3a). We found high coherence values across all unipolar pairs. Coherence values were near unity for adjacent unipolar signals (separated by 25μm) in low frequencies. For these pairs, coherence values decreased with frequency before plateauing at very high values of around 0.8. These very high coherence values in high frequencies in which we would not expect genuine neurophysiological coupling indicate the presence of common signals.

**Figure 3.**
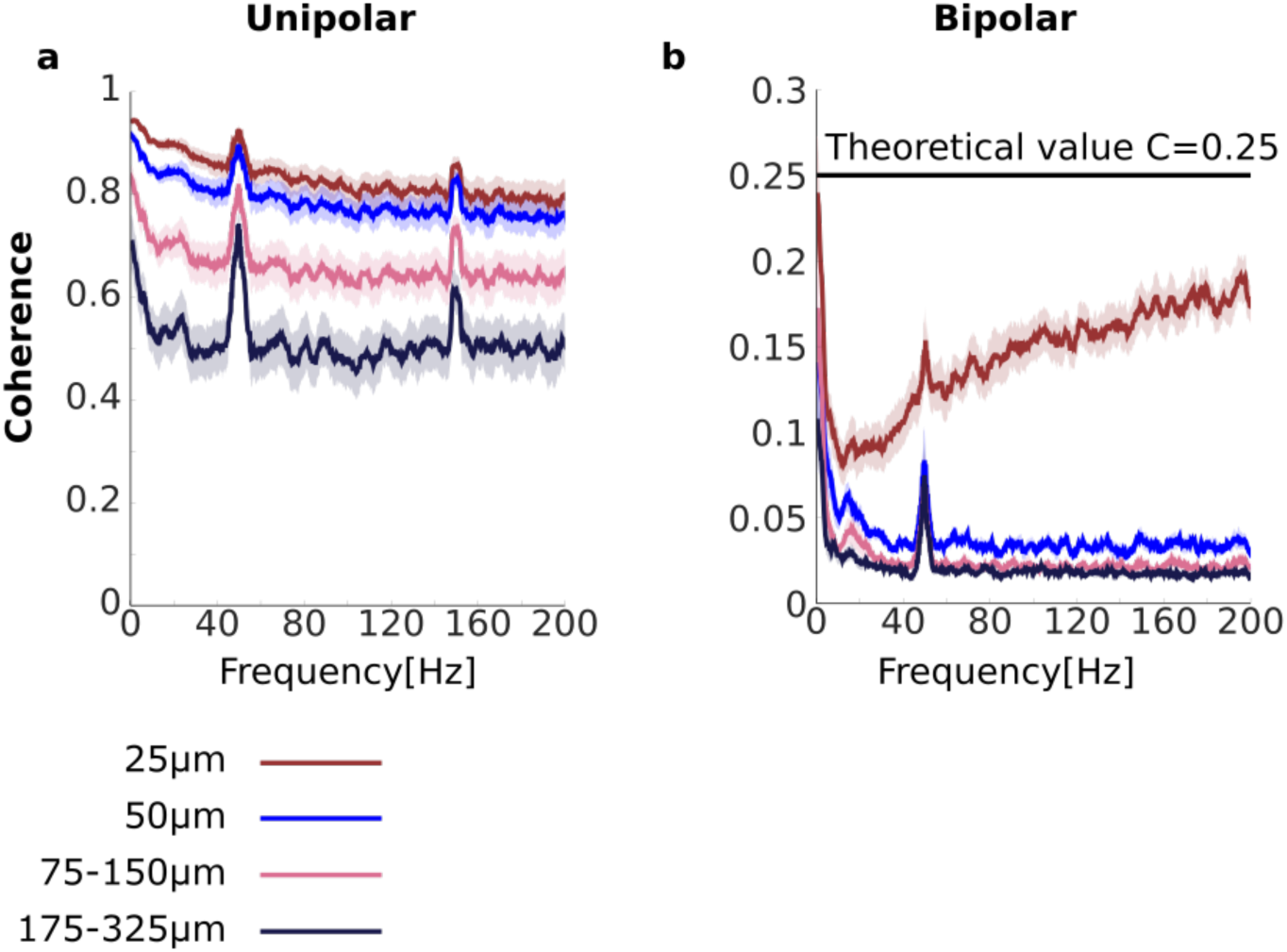
Unipolar and bipolar coherence. **a-b**) Unipolar (**a**) and bipolar (**b**) coherence averaged across signal pairs separated by 25 (brown), 50 (blue), 75-150 (pink), 175-325μm (black) (see Methods for details). The very high unipolar coherences are consistent with the presence of a strong, non-silent reference signal. Bipolar signals that share a unipolar signal in their derivation are separated by 25μm (see Figure 1). Horizontal line at C=0.25 represents theoretical coherence value when the NCR=1. Peaks at 50 and 150Hz are due to line noise and its harmonic. Shaded area represents sem across flies (N=13)

We further found that coherence between unipolar signals decreased with increasing distance between the signals. In low frequencies this reduction may reflect a genuine reduction in neurophysiological coupling with increasing distance. However, the reduction in coherence with distance between the signals was also observed for high frequencies and thus likely to reflect, at least in part, a reduction in the influence of the common signals. This suggests that the properties of the common signals are not constant across all electrodes. For example nearby signals may reflect common signals due to shared neural sources, as well as a common signal due to the non-silent reference.

Unipolar signal pairs that are far apart (175-325μm) are less likely to reflect the activity of a shared neural source than other, closer pairs. However, we found that coherence values were still around 0.5 for these far apart pairs, even for high frequencies for which we would not expect genuine neurophysiological coupling. A likely reason for this high coherence for signals that are far apart is a common signal due to a non-silent reference. Under some further simplifying assumptions we can use the coherence values to estimate the Neural signal to Common signal Ratio (see Eq. (0.14), and Figure 2m). Under these assumptions, coherence values of 0.5 translate to an NCR of approximately 0.41, which means that neural activity is less than half of the magnitude of the common signals. Thus the coherence analysis strongly indicates the presence of a substantial common signal, which we attributed to the non-silent reference electrode located in the flies’ thorax (Figure 1b).

#### 2.3.2 Coherence between bipolar signals is low, as per the theoretical prediction, but coherence between adjacent pairs increases with frequency

Our analyses so far (both power and coherence) strongly indicate that the unipolar signals contain a substantial common signal. According to the mathematical framework, bipolar rereferencing can reduce the effect of common signals on coherence (Section 2.1.2). With bipolar signals, we would expect lower coherence values over all and, in particular, coherence would be near zero for very high frequencies for which we do not expect genuine neurophysiological coupling.

To test this we repeated the coherence analysis with the bipolar signals (Figure 3b). We found that bipolar coherence was indeed much lower than unipolar coherence. For bipolar signals separated by 50-325μm coherence values for low frequencies were in the range 0.025-0.15. Crucially, for higher frequencies we observed near-zero coherence. Taken together these observations suggest that bipolar rereferencing indeed reduces the adverse effects of the common signals. Bipolar coherence also decreased as the distance between the bipolar signals increased, potentially indicating a reduction in neurophysiological coupling with increasing distance.

Surprisingly, coherence between bipolar signals separated by 25μm actually increased with frequency. However, the mathematical framework we provided can fully account for this apparently surprising finding. Bipolar signals separated by 25μm are derived from a shared unipolar signal, whereas bipolar signals separated by 50μm or more are derived from distinct unipolar signals (Figure 1b and c). Coherence between bipolar signals that share a unipolar signal in their derivation constitutes a special case, in which the shared unipolar signal effectively acts as a common signal (Eq. (0.17)). Coherence in the presence of a common signal can be above zero even if the neural signals themselves are independent.

Recall, further, that coherence also increased with frequency in our simulation of a disconnected system in the presence of a common signal (Figure 2l). In that case, we explained it in terms of the analytical relationship between coherence and the NCRs (Figure 2m). We note here that if we assume that the power of the neural signals is equal, then the NCRs equal 1 and coherence equals 0.25 (Eq. (0.13)). As can be seen, coherence approached but never reached this value (Figure 3b, horizontal a black line in). We return to some possible reasons for this in the Discussion.

## 3 Granger causality

### 3.1 Theory

Here we examine how the presence of a reference signal can be incorporated into the Granger causality framework. The mathematical specifics of Granger causality have been discussed extensively (31, 33, 34, 40-44). We cover the relevant details to our analysis in the Methods section. More complete details can be found in the references above.

In the context of our analysis, the key analytical result is the relationship

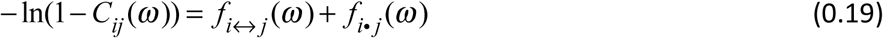

where *f*_*i*↔*j*_(*ω*) represent the sum of Granger causal influences from i to j and j to i, termed total Granger causality, and *f*_*i*·*j*_(*ω*) represents zero-lag or instantaneous effect, termed instantaneous interaction. This result demonstrates that a simple transformation of coherence (left side of (0.19)) can be decomposed into total Granger causality (first term on the right side (0.19)), which capture lagged influences, and instantaneous interaction (second term on the right side of (0.19)), which captures any remaining instantaneous influences, possibly due to exogenous sources (15, 17, 30, 31).

The frequency domain representation of the unipolar signals under the Granger causality framework is

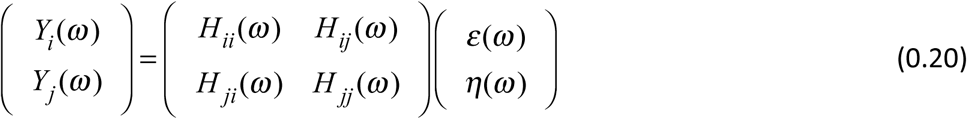

where 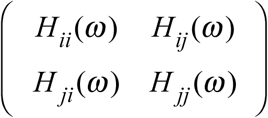 is the transfer function representation of the autoregressive process and 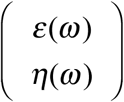 representsthe Fourier transforms of the noise terms (see Methods for details).

Combining the frequency domain representation with the expression of the unipolar signals (0.2) gives

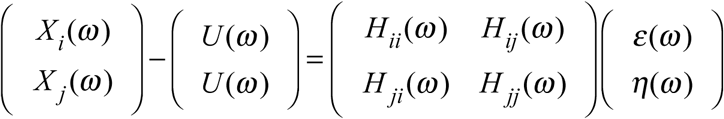

In terms of the neural signals we get

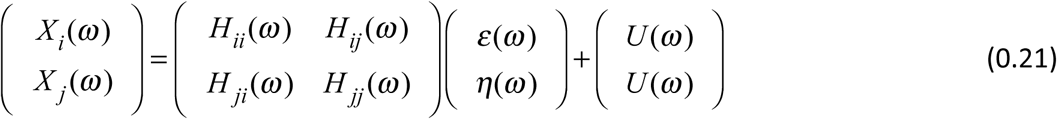

Cast this way we see that the common signal is exogenous to the system. We thus expect that the presence of a common signal will manifest as increased instantaneous interaction.

The same reasoning applies to the case of bipolar signals that share a unipolar signal in their derivation. Substituting the expression for bipolar signals (Eq. (0.7) into (0.20)) we get

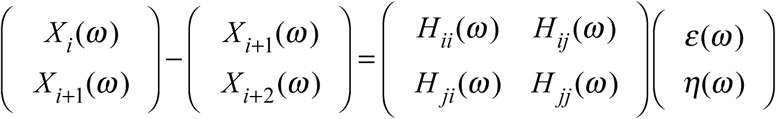

By noting that spectral Granger causality is invariant under scalar multiplication (40) we can use *bY*_*i*+1_(*ω*)→–*bY*_*i*+1_(*ω*) = *X*_*i*+2_(*ω*)– *X*_*i*+1_(*ω*). Rearranging in terms of the neural signals *X*_*i*_(*ω*) and *X*_*i*+2_(*ω*) we get

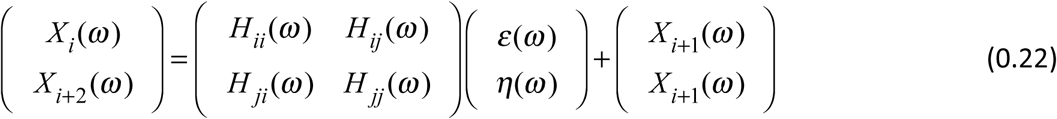

Eq. (0.22) above has the same form as Eq. (0.21). Thus, we expect that instantaneous interaction will be increased for bipolar signals that share a unipolar signal in their derivation.

### 3.2 Example simulations: A common signal increases instantaneous interaction

We next used simple simulations to illustrate how the decomposition in Eq. (0.19) may manifest in empirical analysis. To do this we investigated the same four scenarios as for the coherence analysis in Figure 2; a unidirectionally-connected system and a disconnected system with no common signal, and the same two systems in the presence of an uncorrelated common signal (Figure 4). For each scenario we assessed the quantities in the decomposition in Eq. (0.19); transformed coherence – ln(1 – *C* (*ω*)), total Granger causality (*f*_1_↔2(*ω*)) and instantaneous interaction(*f*_*1·2*_(*ω*)).

**Figure 4.**
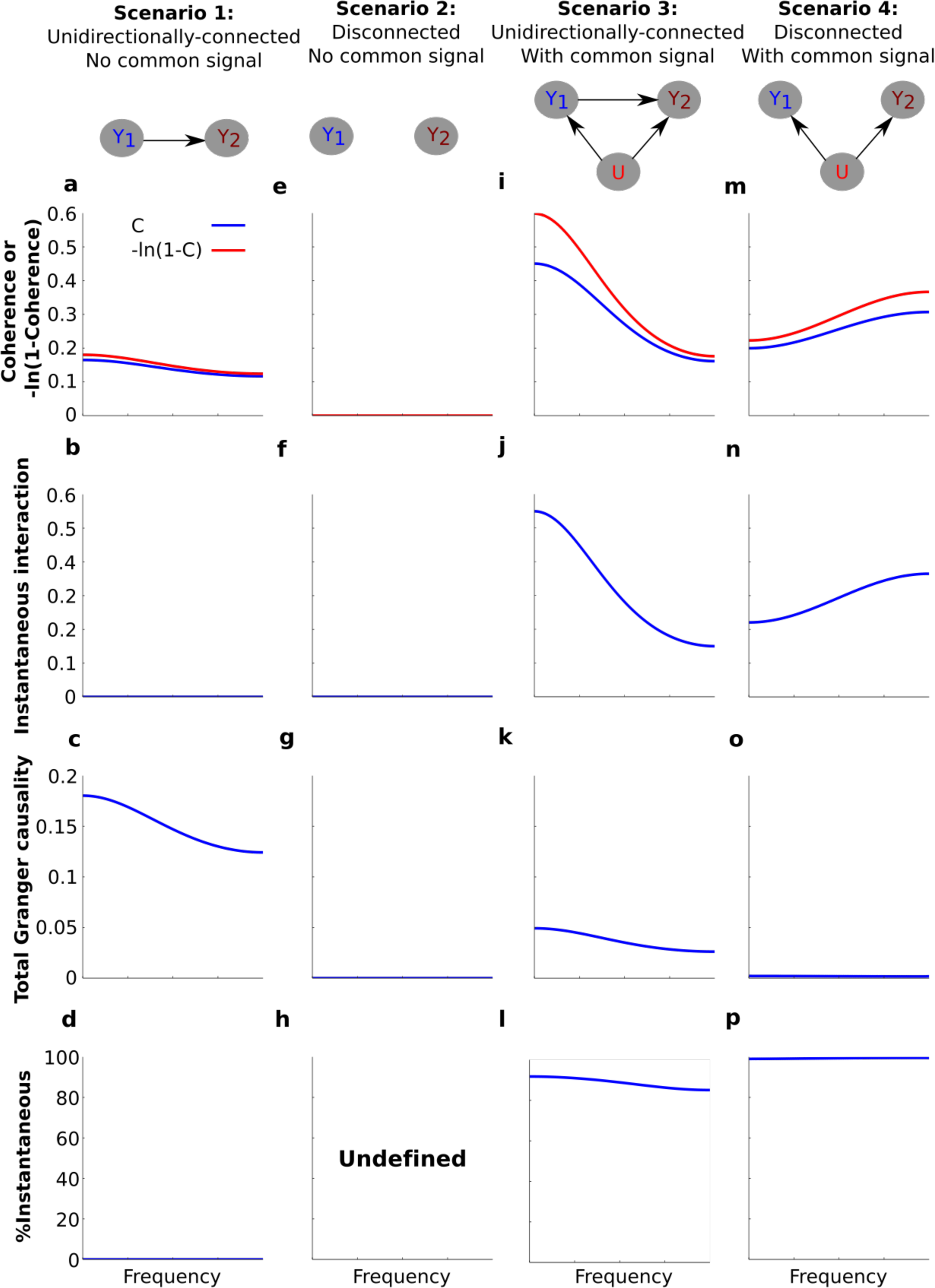
Simulating the effect of a common signal on instantaneous interaction and total Granger causality. Simulation results for **a-d**) the unidirectionally-connected system without a common signal, **e-h**) the disconnected system without a common signal, **i-l**) the unidirectionally-connected system with a common signal, **m-p**) the disconnected system with a common signal. **a,e, i, m**) Transformed coherence (red) and coherence (blue) plotted on the same y-axis scale to facilitate comparison. **b, f, j, n**) Instantaneous interaction. **c, g, k, o**) Total Granger causality. **d, h, l, p**) The percentage of transformed coherence that is accounted for by instantaneous interaction, computed as 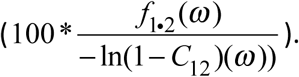. For **h**), the percentage of transformed coherence is undefined due to the division by zero.

For the unidirectionally-connected system (scenario 1) we have already seen that coherence values were low, and decreased with frequency (Figure 2b). In order to decompose this system into total Granger causality and instantaneous interaction we first transformed coherence values into ‐ln(1 – *C*(*ω*)). The transform is unlikely to affect interpretability of the results as it simply ‘stretches’ coherence values. For example coherence values in the [0.01-0.99] range are stretched to the range [0.0101 4.6052]). For the low values of coherence for scenario 1 the effect of the transform is minimal (Figure 4a).

Next we investigated instantaneous interaction. To do this we used the non-parametric Granger causality approach to decompose the spectral density matrix of the system (see Methods). Because the only dependency between the signals for the system is lagged, instantaneous interaction is zero (Figure 4b). Correspondingly, total Granger causality equals transformed coherence for this system (Figure 4c). Because our main interest is in the relationship between instantaneous interaction and coherence, we calculated the percentage of transformed coherence that is accounted for by instantaneous interaction 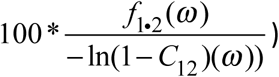 (Figure 4d). Because instantaneous interaction is zero for this system, it accounts for none of the transformed coherence. For the disconnected system without a common signal (Figure 4e-g), coherence, instantaneous interaction and Granger causality are all zero (and % instantaneous interaction is undefined, Figure 4h).

We next investigated how the introduction of a common signal affects the estimation of instantaneous interaction and total Granger causality for the unidirectionally-connected system. We have seen that this results in an overall increase in coherences (Figure 2b and Figure 3i). As we expected, the common signal substantially increased instantaneous interaction (Figure 4j, compared with Figure 4b). Furthermore, introduction of the common signal distorted the estimation of the Granger causality; the total Granger causality was reduced (Figure 4k, compared with Figure 4c). Accordingly, the percentage of instantaneous interaction relative to the transformed coherence is nearly 100%, showing that most of the dependency between the signals is due to instantaneous interaction (Figure 4l).

Finally, we investigated how the common signal affects the estimation of instantaneous interaction and total Granger causality for the disconnected system (Figure 4 m-p). The common signal increased coherence from 0 to 0.3 (Figure 4m, compared with Figure 4e), and magnitude increased with frequency (see also Figure 2l). Instantaneous interaction for this scenario was very high (Figure 4n) and, correspondingly, total Granger causality was very low (Figure 4o). As a result, transformed coherence for this system was almost entirely accounted for by instantaneous interaction (Figure 4p).

In sum, these simple simulations demonstrate how the decomposition in Eq. (0.19) can be applied to the analysis of real data and that a common signal may manifest as increased instantaneous interaction.

### 3.3 Experiment

#### 3.3.1 Instantaneous Interaction accounts for the high coherence observed for the unipolar signals

Our simulations demonstrate that a common signal may manifest as increased instantaneous interaction. If the high coherences we observed for unipolar signals (Figure 3a) are a result of the common signals than we would expect that these would also manifest as increased instantaneous interaction. To investigate this, we decomposed (transformed) coherence into total Granger causality and instantaneous interaction (Figure 5a-d). Due to the transformation, the original unipolar coherence, ranging from ~0.5-0.9 (Figure 3a), were re-scaled to ~0.7-3.0 (Figure 5a). Similar to coherence, transformed coherence decreased with increasing distance between the signals but remained high even for unipolar signals that were far apart (175-325μm). Transformed coherence closely resembled instantaneous interaction in all respects (Figure 5b). Correspondingly, we found that total Granger causality, which captures only lagged effects, was relatively small (Figure 5c). Further, total Granger causality values approached zero as frequency increased, as we would expect if there was no genuine neurophysiological coupling at those frequencies. In sum, this suggests that most of the unipolar coherence was due to instantaneous interaction, which we confirmed by calculating the percentage of transformed coherence that is accounted for by instantaneous interaction (Figure 5d). Together, these findings strongly suggest the presence of common signals in the unipolar data and that this manifests as increased instantaneous interaction.

**Figure 5.**
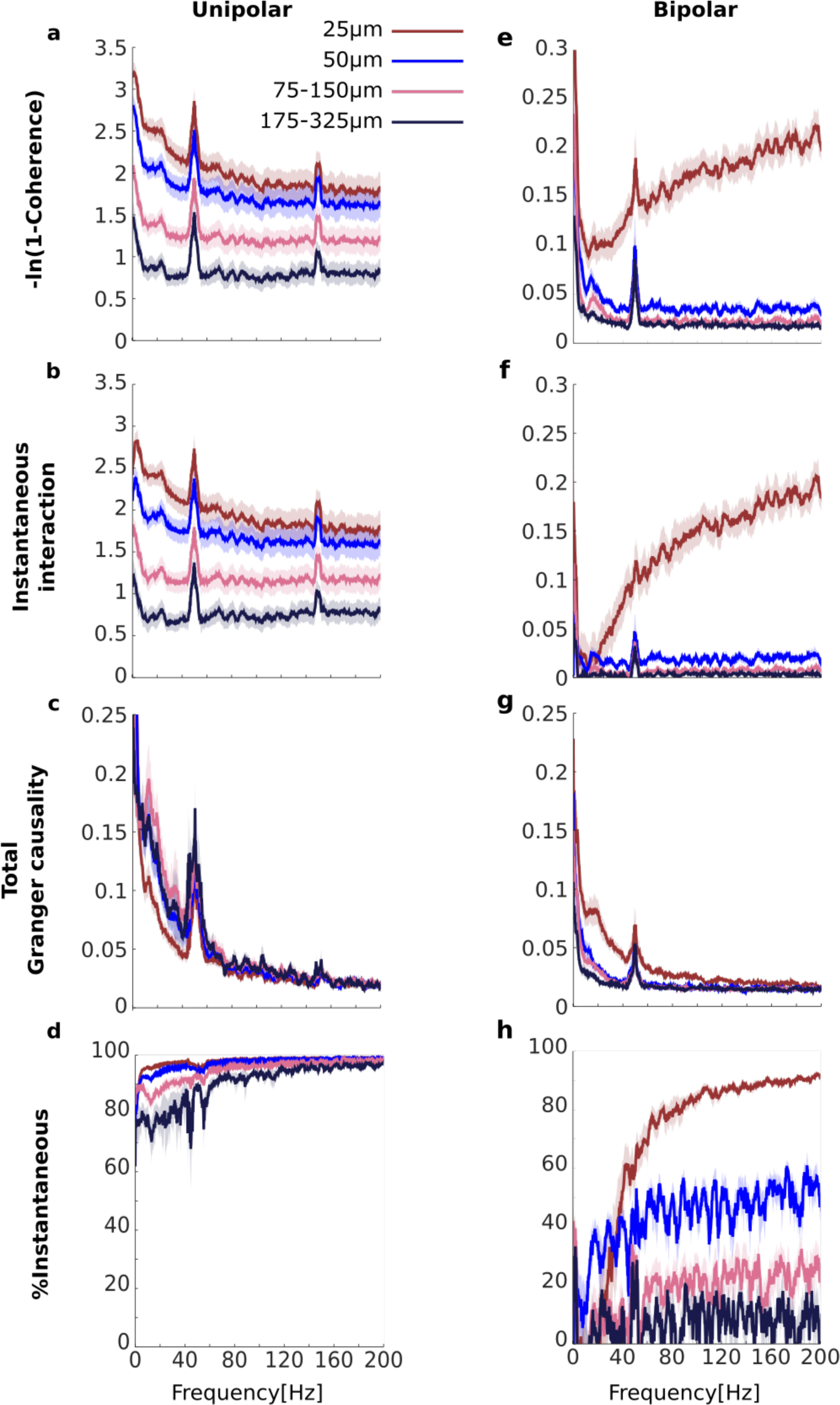
Comparisons of the results for transformed coherence, instantaneous interaction and total Granger causality computed from the experimental data **a-d**) for unipolar and **e-h**) for bipolar signals. **a, e**) Transformed coherence **b, f**), instantaneous interaction, **c, g**), total Granger causality, and **d, h**) the percentage of transformed coherence that is accounted for by instantaneous interaction. The color of each line represents the separation between signal pairs (brown for 25μm, blue for 50μm, pink for 75-150μm and black for 175-325μm, see Methods for details). Pairs separated by 25μm share a unipolar signal in their derivation. Peaks at 50Hz and 150Hz are due to line noise. Shaded area represents sem across flies (N=13).

#### 3.3.2 Instantaneous interaction accounts for the increasing coherence between bipolar signals that share a unipolar signal in their derivation

We have shown that the high unipolar coherences were accounted for by instantaneous interaction, strongly indicating the presence of common signals. Bipolar coherences were generally lower (Figure 3b), which indicates that bipolar rereferencing effectively reduced the effect of the common signals. However, coherence for bipolar signals that shared a unipolar signal in their derivation increased with frequency. We attribute this increase to a common signal due to the shared unipolar signal (Figure 3b). If this common signal is the source of the increasing coherence with frequency, then we would expect this increase in coherence to be accounted for by instantaneous interaction.

We repeated the analysis for the bipolar rereferenced LFP signals (Figure 5e-h). For the bipolar signals, transformed coherence was very similar to the original coherence, including the important characteristic that transformed coherence increased with frequency for bipolar signals that shared a unipolar signal in their derivation (25μm separation show in brown, Figure 5e). Instantaneous interaction (Figure 5f) was much smaller than that for the unipolar signals (Figure 5b), indicating that bipolar rereferencing reduced the effect of the common signals. We also found that instantaneous interaction increased with frequency for bipolar signals that share a unipolar signal in their derivation, but not for pairs derived from independent unipolar signals. This observation is in line with our prediction that this increase was due to the shared unipolar signal. Consistent with this, total Granger causality for these pairs did not increase with frequency, indicating that the increase cannot be attributed to any lagged influences (Figure 5g). Indeed, total Granger causality appeared to decrease with increasing frequency for all bipolar signal pairs. Figure 5h summarizes these results as the percentage of transformed coherence that is accounted for by instantaneous interaction. This clearly showed that the increasing coherence with frequency for bipolar signals that share a unipolar signal in their derivation is dominated by instantaneous interaction. However, even for the bipolar pairs separated by 50μm, which were derived from independent unipolar signals (blue in Figure 5h), instantaneous interaction substantially (~40-60%) contributed to transformed coherence. This could indicate that bipolar rereferencing does not entirely remove the effects of the common signal. We expand on this in the Discussion.

## Discussion

In this paper, we introduced a mathematical framework describing how a common signal influences the estimation of power, coherence, (total) Granger causality and instantaneous interaction. We confirmed our predictions by analyzing local field potentials (LFPs) recorded from fruit flies. The key claims of this paper are five fold. 1) Bipolar rereferencing can increase or decrease the estimated power. Substantially greater unipolar than bipolar power, as observed in our data, is an indication of the presence of common signals (Figure 1d). 2) Theoretically, in the presence of a common signal, unipolar coherence can be greater than zero even if the neural signals are independent (Figure 2). In our data unipolar coherence was high, even for high frequencies for which we would not expect genuine neurophysiological coupling (Figure 3a). 3) Coherence for bipolar signals that share a unipolar signal in their derivation constitutes a special case in which the shared unipolar signal acts as a common signal. Coherence in this case can be above zero even if the neural signals are expected to be independent, as in the high frequency domain of our data (Figure 3b). 4) In general, the effect of a common signal on coherence, (total) Granger causality, and instantaneous interaction will depend on the spectral characteristics of the system and would need to be evaluated on a case-by-case basis. We demonstrate this through simple simulations of a unidirectionally-connected and a disconnected system (Figure 2 and 4). 5) The presence of a common signal may manifest as increased instantaneous interaction as shown in our simulations (Figure 4). In our data instantaneous interaction for the unipolar signals was much higher than that for the bipolar signals (Figure 5). Instantaneous interaction also accounted for the increase in coherence with frequency for bipolar signals that shared a unipolar signal in their derivation. Below we highlight some limitations and future directions.

We believe that our unified theoretical framework serves as a solid foundation for analyzing real neural data. However, a few points should be kept in mind. First, the framework rests on the assumption that the recorded activity can be represented as the sum of ongoing neural activity and a common signal. As such, the framework is suited for guiding the analysis of spontaneous, not evoked, activity, as it does not currently consider the presence of a stimulus. If the only data available corresponds to evoked activity then ‘spontaneous’ activity may be estimated by removing the evoked component (by, for example, averaging across repeated presentations). In such practice, however, care is required to ensure that the resulting data may be reasonably treated as “spontaneous” (45, 46).

Second, in our mathematical treatment we have assumed that the common signal is independent of the neural activity, as this results in relatively simple expressions and is also in line with previous work (e.g. (16, 18-20)). In our simulations (Figure 2 and 4) we were able to satisfy this assumption by construction. However, this may not always be reflective of real data. For example if the reference electrode is placed inside the brain then the recorded reference signal is unlikely to be independent of the neural activity. Even in such a case, the full expressions for power and coherence still hold (e.g. Eqs. (0.3), (0.10) and (0.15)), but inference about any common signals will become more difficult.

Third, for bipolar rereferencing we have assumed that a common signal is identical for sufficiently close electrodes (Eq. (0.6)). This is unlikely to be satisfied *exactly*. As a result, at least some portion of the common signal may still be present in the bipolar signals. Indeed, we found that instantaneous interaction was still present in the bipolar signals, especially for those pairs separated by 50μm (which do not share a unipolar signal in their derivation, blue lines in Figure 5f and h). This may also explain why coherence did not reach its theoretical value for the bipolar signals that shared a unipolar signal in their derivation for high frequencies (Figure 3b).

Lastly, special care is required when interpreting instantaneous interaction. This is because a formal link between instantaneous interaction and a common signal is still unclear. However, we note that some results are available for the related case of measurement noise (47). In addition, concerns about the interpretability of instantaneous interaction have been expressed because it can become negative in certain situations (30, 31, 33). We acknowledge this concern, but also point out that our work here suggests that this quantity may be at least of *empirical* value. For example, it can be used as a diagnostic for the presence of common signals, as we showed here.

We also clarify that even though Granger causality quantifies lagged effects, its estimation in the presence of a common signal maybe misleading (15, 17, 47, 48). Indeed, in our simulations the presence of a common signal resulted in lower Granger causality values (Figure 4k) than the ground truth without a common signal (Figure 4c). With our analysis of the unipolar data, total Granger causality in lower frequencies (<50Hz) for adjacent pairs (separated by 25μm) were actually smaller than those for pairs that were separated by more than 50μm, which is physiologically highly questionable (Figure 5c). With our analysis of the bipolar data, however, higher total Granger causality for closer pairs were observed in low frequency range (<50Hz, Figure 5g), which is much more physiologically plausible.

## Conclusion

In this work we focused on detecting and assessing common signals in multi-electrode neurophysiological recordings. To do this, we contrasted in depth analysis of the unipolar and bipolar signals with the latter likely to reduce the effect of common signals present in the former. We did not consider numerous other techniques for removing common signals, such as other referencing techniques (e.g. (16, 18, 19)), methods based on Independent or Principal Component analysis (e.g. (20, 22, 23) or other linear decompositions (24-26)), or methods based on more detailed modeling of the dynamics (e.g. (47-50)). We did not assess the many other functional and effective connectivity techniques (e.g. (51-53)). Instead, we focused on bipolar rereferencing together with coherence and Granger causality metrics, which are all widely adopted. In addition coherence and Granger causality can be analytically related in a (relatively) straightforward manner. By exploiting analytic relationships between other connectivity metrics it may be possible to gain an even deeper understanding into the nature of any common signals. Our approach combined with other methods for assessing common signals (54) will reduce the chance of misinterpreting functional and effective connectivity analysis, the key analysis techniques in modern systems and cognitive neuroscience (1-5, 7-9).

## Methods

In the main text, we have provided the core theoretical background, simulation, and analysis of the experimental data. Here, we provide further methodological details. We first cover the mathematical formulation of spectral Granger causality. We then provide full details of the simulations, followed by complete details of the LFPs data analysis.

## Spectral Granger causality

In this section we briefly recap the theoretical background for spectral Granger causality. Here we focus on the lesser-known relationship between coherence, Granger causality and instantaneous interaction and we only rehearse the relevant components to our analysis, generally following the treatment of (30, 31). More complete details can be found in (31, 33,34,40-44).

In simple terms, a signal *y*_*i*_ is said to *Granger-cause* a signal *y*_*j*_ if past values of *y*_*i*_ improve predictions of future values of *y*_*j*_. This notion is quantified using the framework of autoregressive processes.

Consider two stationary time series represented by the standard autoregressive process

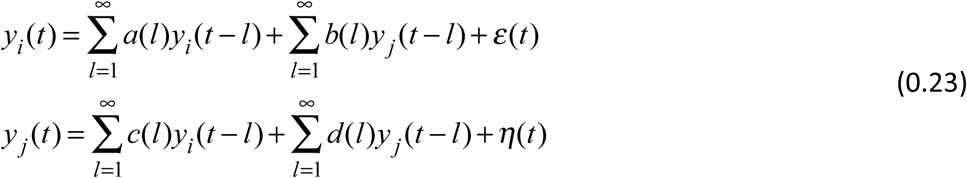

where *a*,*b*,*c* and *d* represent the autoregressive coefficients and the index *l* represents the lag. *ε*(*t*) and *η*(*t*) are zero-mean Gaussian noise sources with covariance matrix given by

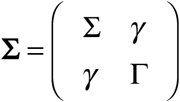

where var(*ε*(*t*)) = ∑,var(*η*(*t*)) = Γ and *cov*(*ε*(*t*),*η*(*t*)) = *γ*.

To obtain the spectral formulation of Granger causality we first express the autoregressive process in the frequency domain. By introducing the lag operator *L*, *Ly*_*i*_ (*t*) = *y*_*i*_ (*t* – 1) we can rewrite Eq. (0.23) in matrix form as

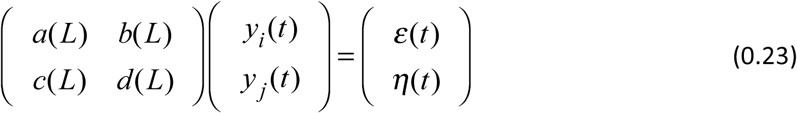

where *a*(0) = 1,*b*(0) = 0, *c*(0) = 0, *d*(0) = 1. Taking the Fourier transform of both sides gives

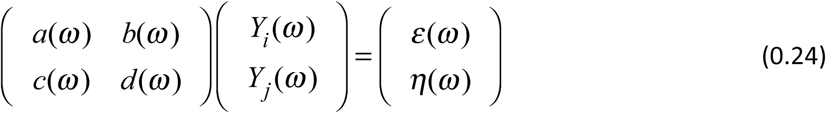

By using 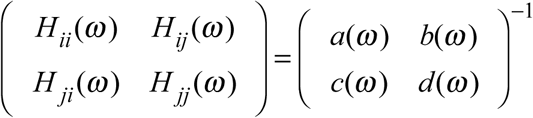 we can rewrite Eq.(0.24) in transfer function format as

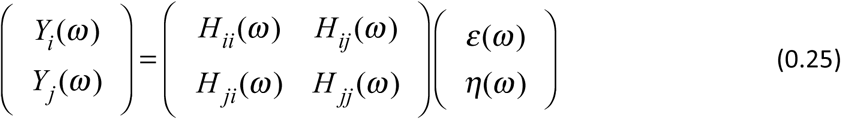

The spectral density matrix is obtained by multiplying both sides by the conjugate transpose of each side, denoted by ^*^, yielding

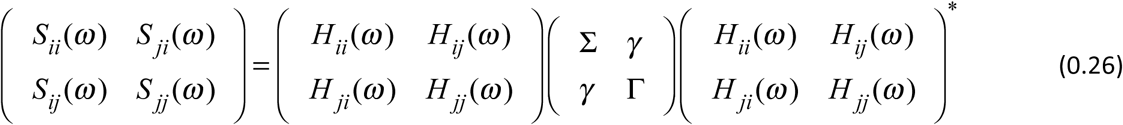

This can be compactly written in matrix form as

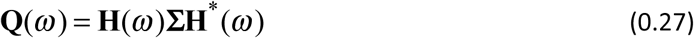

where 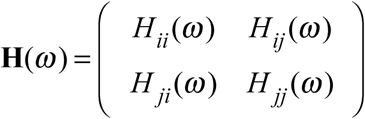 and 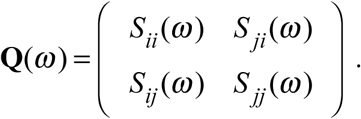

The diagonal elements of the spectral density matrix correspond to the auto-spectra and the off diagonal elements correspond to the cross-spectra (31). Note that coherence can be directly calculated from the spectral matrix (Eq. (0.9)).

Spectral measures of total interdependence *f*_*ii*_(*ω*), GC influence from Yi to Yj *f*_*i*_→*j*.(*ω*) and GC influence from Yj to Yi *f*_*i*←*j*_(*ω*) are defined as

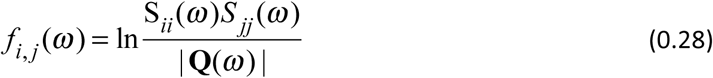

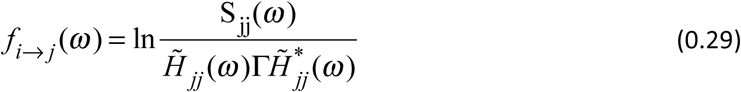

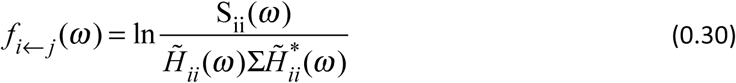

where *H̃*_*jj*_(*ω*) = *H*_*jj*_(*ω*) + (*γ* / Γ)*H*_*ij*_(*ω*) and *H̃*(*ω*) = *H*_*ii*_(*ω*) + (*γ* / ∑)*H*_*ij*_(*ω*). These transformations are irrelevant if *γ* = 0.

A simple transform relates coherence and total interdependence

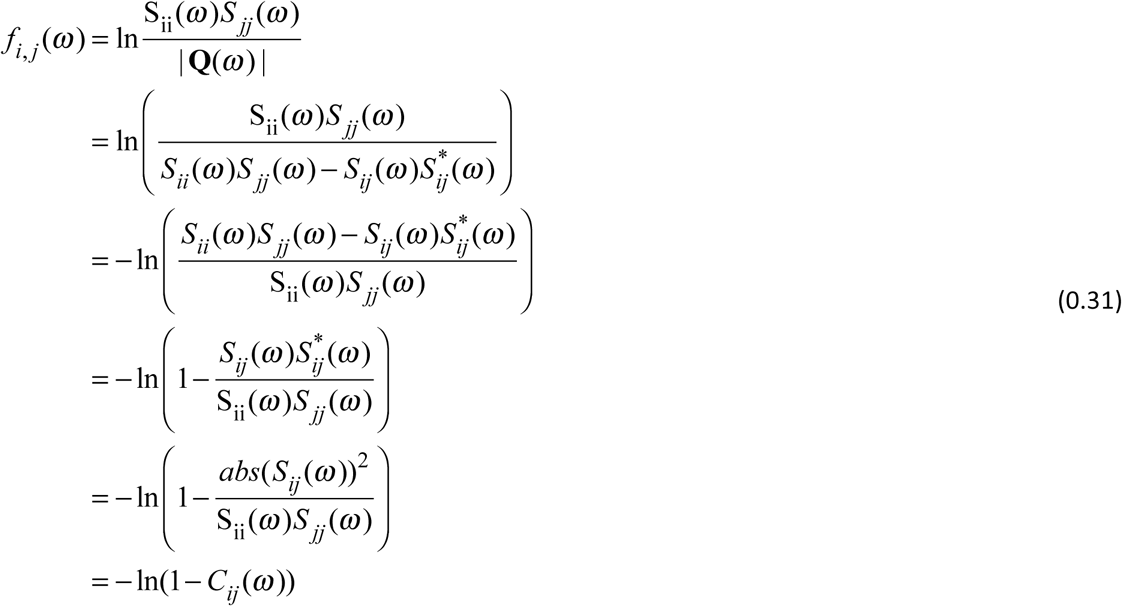

By subtracting *f*_*i*_→*j*(*ω*) and *f*_*i*_←*j*(*ω*) from *f*_*i*,*j*_(*ω*) we can get what is known as ‘instantaneous causality’ (*f*_*i*·*j*_(*ω*) (30, 31))

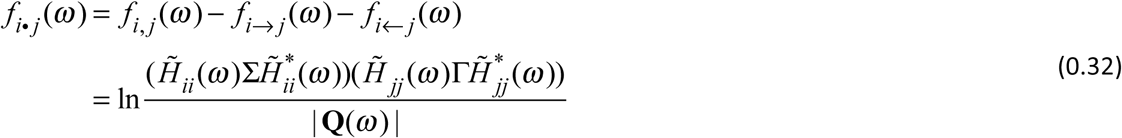

However, we feel that the term *instantaneous causality* is confusing in so far as (Granger) *causality* reflects *time-lagged* influences. For this reason we refer to this term as *instantaneous interaction* throughout the manuscript.

Using instantaneous interaction and the Granger causal influences we obtain the following decomposition of total interdependence

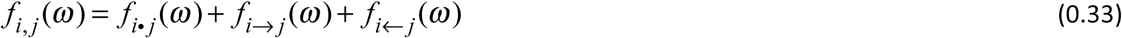

The relationship between coherence, Granger causality and instantaneous interaction is given by

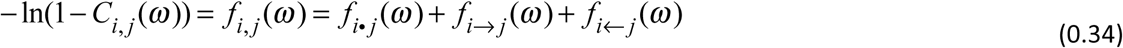

If we disregard the directionality of the influence, we can simply contrast ‘lagged’ (*f*_*i*→*j*_(*ω*) + *f*_*i*←*j*_(*ω*) and ‘instantaneous’ (*f*_*i*·*j*_(*ω*)) influences. To do this we introduce *total Granger causality f*_*i*↔*j*_(*ω*)

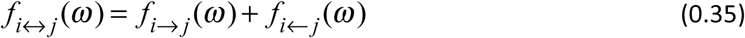

We thus rewrite Eq. (0.34) as

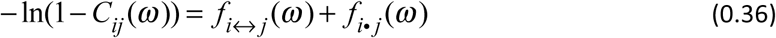

This equation demonstrates that under the autoregressive framework of Granger causality, (a transformation of) coherence (left side of (0.36)) can be decomposed into total Granger causality (first term on the right side (0.36)), which capture lagged influences, and instantaneous interaction (second term on the right side of (0.36)), which captures any remaining instantaneous influences, possibly due to exogenous sources. This relationship forms the basis of our simulation examples and experimental data analysis.

## Non-parametric estimation of spectral Granger causality

Spectral Granger causality analysis can be carried out either parametrically or non-parametrically. In the parametric approach the autoregressive model in Eq. (0.23) is first fit to the data. Once the parameters of the models have been estimated Eqs. (0.23) – (0.27) are used to obtain the transfer function **H**(*ω*), noise covariance matrix **∑** and spectral density matrix **Q**(*ω*). Using these quantities *f*_*i*→*j*_(*ω*), *f*_*i*←*j*_(*ω*) and *f*_*i*·*j*_(*ω*) can be calculated as per Eqs. (0.29), (0.30) and (0.32).

In the non-parametric approach proposed in (34), **H**(*ω*) and **∑** are obtained directly by factorizing the spectral density matrix **Q**(*ω*). This corresponds to obtaining the right side of Eq. (0.27) directly from its left using a factorization procedure described by (55), without explicitly fitting the autoregressive model. Thus, non-parametric estimation is entirely dependent on the estimation of the spectral density matrix. One advantage of non-parametric over parametric estimation of GC influences is that it does not require specification of the autoregressive model order (34).

Recall that coherence (Eq. (0.9)) is also directly estimated from the spectral density matrix Thus, once the spectral density matrix has been estimated, power, coherence, total Granger causality and instantaneous interaction can all be derived. For our simulations the spectral density matrix is obtained analytically while for the experimental sections the spectral density matrix is estimated empirically. We provide further details on these below.

## Simulations

In our simulations we explored how coherence, total Granger causality and instantaneous interaction are affected by the presence of a common signal. To do this we explored four scenarios. The first two scenarios correspond to a unidirectionally-connected and a disconnected (meaning that the cross-spectrum between the signals is zero for all frequencies) system. These two scenarios establish the ‘ground truth’ for coherence, total Granger causality and instantaneous interaction in the absence of a common signal. The remaining two scenarios examine how these quantities are affected by a common signal. Following the notation of the unipolar signals the four scenarios can be described as

|  |  |  |
| --- | --- | --- |
| | Unidirectionally-connected<br>$(X_1'(\omega)X_2'^*(\omega) \neq 0)$ | Disconnected<br>$(X_1'(\omega)X_2'^*(\omega) = 0)$ |
| No common signal | Scenario 1<br>$y_1(t) = x_1(t)$<br>$y_2(t) = x_2(t)$ | Scenario 2<br>$y_1'(t) = x_1'(t)$<br>$y_2'(t) = x_2'(t)$ |
| With common signal | Scenario 3<br>$y_1(t) = x_1(t) + u(t)$<br>$y_2(t) = x_2(t) + u(t)$ | Scenario 4<br>$y_1'(t) = x_1'(t) + u(t)$<br>$y_2'(t) = x_2'(t) + u(t)$ |

We clarify that these scenarios are only indirectly related to the choice of referencing scheme. Instead, these simulations directly investigate the effects of a common signal. This can correspond to either unipolar signals which are affected by a non-silent reference signal, or to bipolar signals which share a unipolar signal in their derivation.

## Simulation framework

For our simulations we used a=0.1, b=0.4 and d=0.1 in Eq. (0.18). These parameters settings result in power and coherence spectra that are roughly reflective of biological systems (Figure 2e,i). The noise terms were chosen as uncorrelated noise sources with unit standard deviation.

When modeling the disconnected system we ensured that the power spectra equaled the power spectra of the unidirectionally-connected system. That is, we set
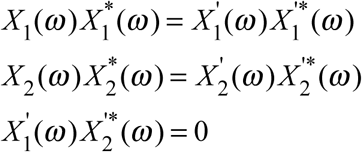

We aimed to do this because coherence, total Granger causality and instantaneous interaction depend on the power spectra of the signals (see Eq. (0.9) for coherence and Eq. (0.29) for Granger causality). By equating the power we ensured that the two systems are not distinguishable based on the power alone. Note that simply setting the autoregressive coefficient b=0 in Eq. (0.18) would ensure the system is disconnected but would also change the power spectrum of *X*_1_(*ω*)

We modeled the common signal for scenarios 3 and 4 as an uncorrelated white noise signal,*u*(*t*). We set the power of the signal (*U*(*ω*)*U*^*^ (*ω*)) to the mean power of the neural signals across frequencies

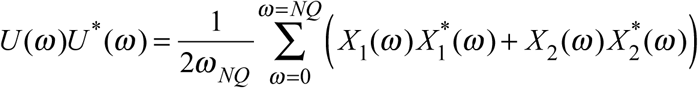

where *ω*_*NQ*_ corresponds to the Nyquist frequency which is arbitrary for our simulation.

## Simulations for the unidirectionally-connected system (Scenarios 1 and 3)

As mentioned above, the key quantity to estimate is the spectral density matrix, from which power, coherence, total Granger causality and instantaneous interaction can all be derived.

For the unidirectionally-connected system with no common signal (scenario 1) one can use the autoregressive description of the system and Eqs. (0.23) – (0.27) to obtain the spectral density matrix 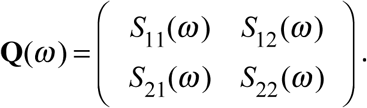. The spectral density matrix of an autoregressive process can be obtained using the MATLAB (MathWorks) function *ft_freqanalysis.m* function from the FieldTrip toolbox (56). Coherence and instantaneous interaction are obtained using the FieldT rip function *ft_connectivityanalysis.m.* Total Granger causality is obtained by first assessing the Granger causal influences from y1 to y2 and from y2 and y1 (also using *ft_connectivityanalysis.m*) and summing them as per Eq. (0.35).

Eqs. (0.4) and (0.11) describe the effect of the common signal (scenario 3) on power and coherence respectively. Unlike power and coherence, there are currently no analytical results describing how a common signal affects instantaneous interaction and total Granger causality. However, we note that if we know the effect of a common signal on the spectral density matrix, then we can use the non-parametric estimation procedure to obtain the affected transfer function and noise covariance matrix

To see the effect of the common signal on the spectral density matrix we re-write scenario 3 in the frequency domain as

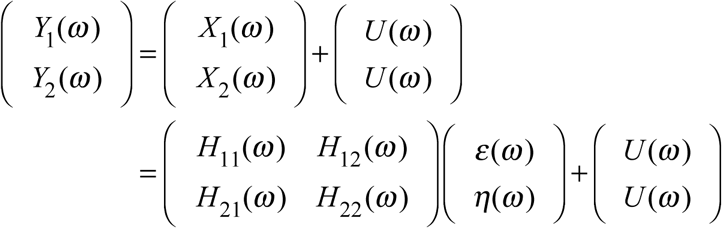

Next we can calculate the spectral density matrix by multiplying both side by the conjugate transpose

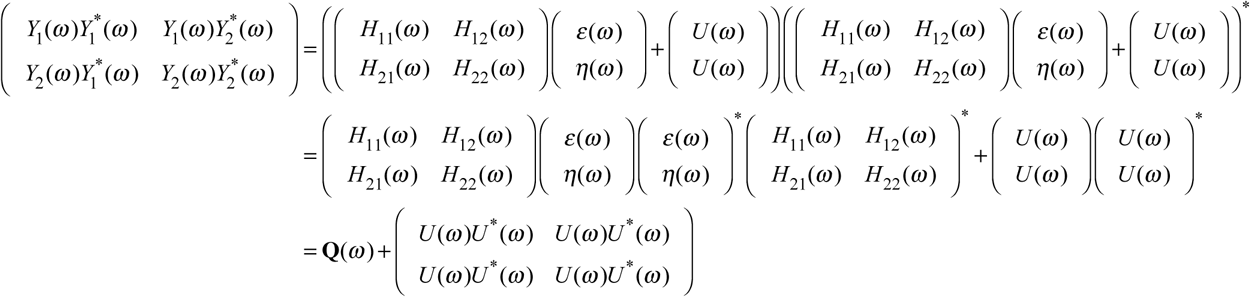

Note that the cross-spectra between the common signal (*U*(*ω*)) and any other quantity (e.g.,*H*(*ω*),*ε*(*ω*),*η*(*ω*)) vanish since we assumed independence between them. Thus, to obtain the spectral density matrix (**Q**^*c*^ (*ω*)) of the unidirectionally-connected system with common signal, we simply add the power of the common signal (*U*(*ω*)*U*^*^(*ω*) to the spectral density matrix of the unidirectionally-connected system without common signal (**Q**(*ω*)). Once the affected spectral density matrix is known, total Granger causality and instantaneous interaction are derived as before.

## Simulation for the disconnected system (Scenarios 2 and 4)

For the disconnected system (scenario 2) we followed a similar procedure but used a modified spectral density matrix 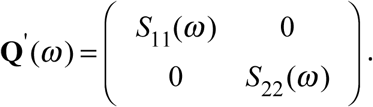.This spectral density matrix describes a disconnected system with identical power spectra to the unidirectionally-connected system. Coherence, instantaneous interaction and total Granger causality are all zero for the disconnected system when there is no common signal.

The effect of the common signal (scenario 4) on the power spectra is identical to the unidirectionally-connected system (Eq. (0.4)). However the effect on coherence is now given by Eq. (0.12). Further, we can estimate the Neural signal to Common signal Ratio (NCR) from coherence using Eq. (0.14).

The effect of the common signal on instantaneous interaction and total Granger causality is obtained as for the unidirectionally-connected system, but using **Q**’ (*ω*) instead of *Q*(*ω*).

We note that an alternative simulation approach would be to use the auto-regressive descriptions to generate time domain data with or without a common signal, and then analyze coherence, total Granger causality and instantaneous interaction in these data (e.g. (30, 31)). If a large enough amount of data is used then the two approaches yield identical results (which we also verified in simulation). However, simulating time domain data involves more parameters. At the very least we would need to decide on the amount of data (number of trials/length of trials/sampling rate). We would also need to describe the frequency domain analysis, which can include parameters such as the number of tapers for multitaper analysis. Another drawback is that these simulations will suffer from empirical biases when using finite data, which are known to affect coherence (57) and Granger causality (58). While all of these are important aspects that deserve further consideration, we feel that they would unnecessarily complicate our simulation procedure. They may also make interpretation more difficult, as differences between the scenarios may dependent on the choice of these additional parameters.

Matlab code for the simulations is available from *https://github.com/DrorGitHub/A-unified-framework-for-dissecting_simulations*

## Experimental setup

To test the utility of the mathematical framework we analyzed previously published data of spontaneous LFPs recorded from the brains of fruit flies. The full experimental details can be found in ((36, 59)). We briefly recap the relevant details below.

Thirteen female *D. melanogaster* flies were tethered and positioned on an air-supported Styrofoam ball. Linear silicon probes with 16 electrodes separated by 25μm were inserted laterally to the eye of the fly until the most peripheral electrode site was just outside the eye. This probe covers approximately half of the brain (Figure 1a). A fine tungsten wire was inserted in the thorax and used as a reference electrode (Figure 1b).

## Data analysis

Data was recorded at 25 kHz and downsampled to 1000Hz and the most peripheral electrode site was removed from the analysis. We analyzed both unipolar and bipolar signals. Unipolar signals correspond to the remaining 15 channels that were recorded with reference to the thorax (Figure 1b). The bipolar signals were obtained by subtracting adjacent unipolar signals, providing another set of 14 signals (Figure 1c).

Because we use a linear array recording, the separation between electrodes increases linearly in multiples of 25μm (Figure 1a). We take the position of the unipolar signal to be the position of the electrode. We take the position of the bipolar signal to be the mid point of the two unipolar signals used in its derivation. Thus, the separation between both unipolar and bipolar signals increases in multiples of 25μm. Adjacent bipolar signals are separated by 25μm and share a unipolar signal in their derivation, whereas bipolar signals separated by 50μm or more are derived from four distinct unipolar channels (Figure 1c).

We analyzed 8 consecutive epochs of 2.25s corresponding to 18s of spontaneous LFPs (as in (59)). We removed line noise from each epoch and signal (unipolar or bipolar) using the rmlinesmovingwinc.m function from the Chronux toolbox (http://chronux.org/; Mitra and Bokil, 2007) with three tapers, a window size of 0.75s and a step size of 0.375s. We preprocessed each epoch and signal by linearly detrending, followed by subtraction of the mean and division by the standard deviation across time points (i.e., z-scoring across time). Example of the resulting pre-processed unipolar and bipolar LFPs from one fly are shown in Figure 1b and c respectively.

### Spectral density matrix estimation

We computed the auto‐ and cross-spectra, 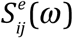, for each unipolar signal pair *Y*_*i*_(*t*) and *y*_*j*_(*t*) (i,j =[1-15]) for each epoch *e* (1-8) over the 2.25s using the multitaper method based on the MATLAB Chronux toolbox (http://chronuxorg/; (60)) function mtspectrumc.m with 9 tapers, giving a half bandwidth of 2.22Hz (61). We obtained the spectral density matrix 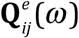 for signals *Y*_*i*_(*t*) and *Y*_*i*_(*t*) by setting the diagonal elements to the auto-spectra and the cross-diagonal elements to the cross-spectra (31)

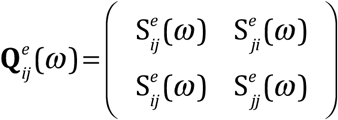

To provide one estimate of the unipolar spectral density matrix **Q**_*ij*_(*ω*), we averaged 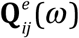 across the 8 epochs (e=1…8). To estimate the bipolar spectral density matrices *b***Q**_*ij*_(*ω*), we repeated this for the bipolar signals. Power, coherence, total Granger causality and instantaneous interaction were estimated from the unipolar and bipolar spectral density matrices, as described below.

### Power analysis

We obtained an overall estimate of unipolar signal power by averaging the power spectra across all channels in units of 10log10

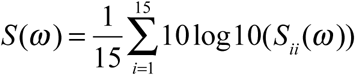

We obtained analogous estimates of bipolar power *bS*(*ω*) using

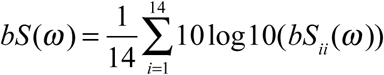

The group average unipolar and bipolar signals power is reported in Figure 1d.

### Coherence analysis

Coherence (Eq. (0.9)) is directly calculated from the cross-spectral density matrices. As discussed extensively in the main text, the effect of a common signal on coherence may depend on the distance separating the two signals. To assess any effects due to the distance, we grouped channel into 4 groups: long-(175-325μm), mid-(75-150μm), short-distance (50μm), and adjacent pairs (25μm).

Specifically, for the unipolar signals we report

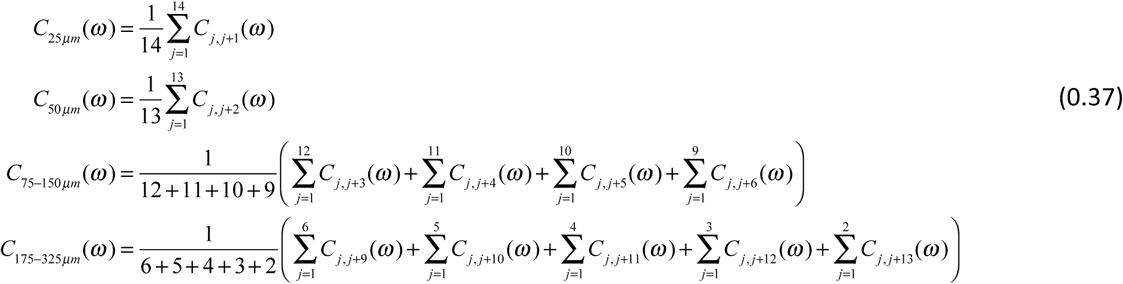

Equivalently, for bipolar coherence we report

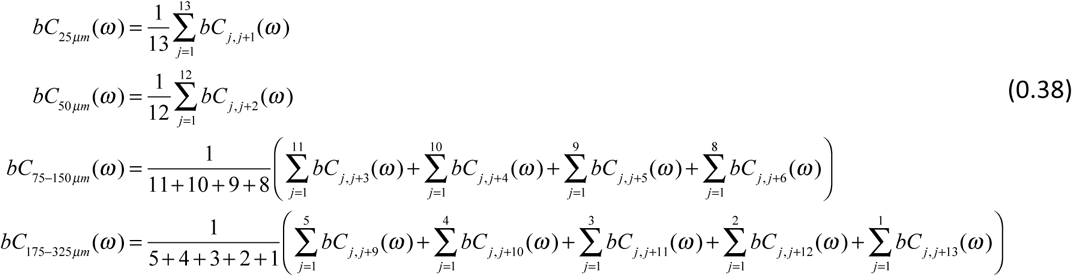

Examining finer channel groupings did not reveal any additional insights.

### Granger causality analysis

To calculate total granger causality and instantaneous interaction we factorized the unipolar (**Q**_*ij*_(*ω*)) and bipolar (*b***Q**_*ij*_(*ω*)) spectral density matrices as described in *Non-parametric estimation of spectral Granger causality*. We averaged total granger causality and instantaneous interaction per channel separation in the same way we did for coherence (Eqs. (0.37) –(0.38))

